# Alfalfa leaf curl virus is efficiently acquired by its aphid vector *Aphis craccivora* but inefficiently transmitted

**DOI:** 10.1101/2020.01.15.908046

**Authors:** Faustine Ryckebusch, Michel Peterschmitt, Martine Granier, Nicolas Sauvion

## Abstract

Alfalfa leaf curl virus (ALCV) is the first geminivirus for which an aphid transmission was reported. Transmission by *Aphis craccivora* was determined previously to be highly specific and circulative. Using various complementary techniques, the transmission journey of ALCV was monitored from its uptake in an infected plant tissue up to the head of its vector. ALCV was shown to be restricted in the phloem using fluorescent in situ hybridization (FISH) and electropenetrography (EPG) monitoring of virus acquisition. Furthermore, the virus is heterogeneously distributed in the phloem as revealed by FISH and qPCR quantification of the viral DNA acquired by aphids monitored by EPG. In spite of the efficient ingestion of viral DNA, about 10^6^ in a 15-hour feeding on ALCV infected plants, the individual transmission rate was at a maximum of 12%. Transmission success was related to a critical viral accumulation, around 1.6×10^7^ viral DNA copies per insect, a threshold that needs generally more than 48 hours to be reached. Moreover, whereas the amount of acquired virus does not decrease over time in the whole aphid body, it decreased in hemolymph and heads. ALCV was not detected in progenies of viruliferous aphids and had no effect on aphid fitness. Compared to geminiviruses transmitted by whiteflies or leafhoppers or to luteovirus transmitted by aphids, the transmission efficiency of ALCV by *A. craccivora* is low. This result is discussed in relation to the aphid vector of this geminivirus and the agroecological features of alfalfa, a hardy perennial host plant.

## INTRODUCTION

Generally, plant viruses belonging to the same family are transmitted according to the same transmission mode (1, 2). In particular, the viruses of the family *Geminiviridae* are all known or suspected to be transmitted according to the circulative mode, although they are transmitted by vectors belonging to different hemipteran groups. Thus, viruses of the genus *Begomovirus* are transmitted by whiteflies (3), those of the genera *Mastrevirus, Curtovirus, Becurtovirus, Turncurtovirus* and probably *Eragrovirus* are transmitted by leafhoppers (4-6), those of the genera *Topocuvirus* and *Grablovirus* are transmitted by treehoppers (7), and those of the genus *Capulavirus* are transmitted by aphids (8-10). This means that geminiviruses were able to adapt to different insect environment and particularly to counteract defense mechanisms and achieve the complex virus-vector interactions associated with circulative transmission. Thereon, compared to *Luteoviridae* or *Nanoviridae*, two families of viruses also transmitted in a circulative manner but quasi exclusively by aphids (11), the family *Geminiviridae* stands in striking contrast. With the exception of the recently discovered whitefly transmitted polerovirus (12), it seems that members of *Luteoviridae* or *Nanoviridae* were not able to evolve all the necessary interactions to be transmitted by other hemipteran groups, which altogether suggests that the adaptation to other hemipteran groups is not straightforward.

Surprisingly, whilst diseases caused by leafhopper and whitefly transmitted geminiviruses were reported more than 100 years ago (13-16), the aphid transmitted geminiviruses were discovered less than 10 years ago (17). The circulative leafhopper transmission of maize streak virus (MSV), the type member of the genus *Mastrevirus* and that of beet curly top virus (BCTV), the type of the genus *Curtovirus* were described in the 30s using laborious infectivity based techniques (18, 19). It was also in the 30s that the transmission of a geminivirus by the whitefly *Bemisia tabaci* was described for the first time with the cotton leaf curl virus (CLCV) (20), a member of the genus *Begomovirus*.

Several reasons may explain the much later detection of viruses classified in the genus *Capulavirus*. Firstly, two of them, euphorbia caput-medusae latent virus (EcmLV) and plantago lanceolata latent virus (PlLV) (17), did not attract attention because they are latent and infect non-cultivated hosts. Secondly, although french bean severe leaf curl virus (FbSLCV) was isolated from a cultivated host (17), its economic importance was relatively low because its incidence is generally below 2% (Akram, personal communication). Thirdly, the leaf curl symptoms associated with alfalfa leaf curl virus (ALCV) have been previously reported to be induced by a rhabdovirus (21, 22). Although it is presently not known how much ALCV contributes to the leaf curl disease of alfalfa, overall the results indicate that so far, the dissemination and economic importance of capulaviruses in cultivated plants are low.

Representatives of the genus *Begomovirus, Mastrevirus* or *Curtovirus* are among the most economically important plant viruses globally, and the genus *Begomovirus* is the largest genus of plant viruses (23, 24). Thus, while Aphididae is the family of the order Hemiptera that contains most of the plant viruses vector’s (11), it is presently not known why aphid-transmitted geminiviruses were much less invasive in cultivated plants than leafhopper and whitefly-transmitted geminiviruses. A tantalizing hypothesis is that the biological specificity of aphids (for example specific structure of the virus-crossed organs or insect-specific defense mechanisms) could not lead to a high transmission efficiency of geminiviruses.

To test the hypothesis of low transmission efficiency of capulaviruses, we monitored the transmission pathway of ALCV from the infected plant tissues to the salivary glands of its vector *Aphis craccivora* using complementary techniques to localize and quantify the virus in the different phases of its transmission journey. While ALCV acquisition, accumulation and persistence are as efficient as those reported with other circulative viruses, the transmission rate is much lower than that reported with non-aphid transmitted geminiviruses.

## METHODS

### Plant and insect material

All transmission tests were conducted using broad bean plants (*Vicia faba* L. cv. ‘Séville’). Fluorescent in situ hybridization (FISH) analysis was performed on both broad bean and *Nicotiana benthamiana* plants. Plants were grown in a P2 containment chamber under 16h light at 26 °C, and 8h dark at 24°C.

Individuals of *A. craccivora* Koch, 1854, were collected in 2015 by G. Labonne (INRA, France) on *Robinia pseudoacacia* L. (False acacia) near Montpellier (France). A rearing was initiated and maintained on broad bean plants (*Vicia faba* L. cv. ‘Séville’) under 16h light at 24°C and 8h dark at 21°C.

### Preparation of agroinfectious clones and agroinoculation

The agroinfectious clone of ALCV was reported previously (8). Agrobacteria were cultured and inoculated to broad bean plants as described Ryckebusch *et al.* (9).

### Plant DNA extraction and detection of ALCV DNA by PCR

ALCV infection of broad bean plants was monitored 4 to 6 weeks after inoculation by symptom observation and/or by PCR-mediated detection of viral DNA in total plant DNA extracts as described Ryckebusch *et al.* (9).

### Mechanical inoculation of ALCV

One gram of leaf material was collected from broad bean plants 4 weeks after their agroinoculation with ALCV. According to a first extraction method, the plant material was ground in an ice-cold mortar with 4 ml of a cold 0.03M sodium phosphate buffer (Na_2_HPO_4_) containing 0.2% sodium diethyldithiocarbamate (DIECA). After the addition of 0.2 g activated charcoal and 0.4 g carborundum, the plant extracts were rubbed onto the upper side of the youngest leaves of twenty-two 10 day-old broad bean plants. Plants were abundantly rinsed with water 5 min after their inoculation.

A second trial was performed as described by Susi *et al.* (10): two grams of leaf material were collected from broad bean plants 33 days after their agroinoculation with ALCV. Plant material was ground in a cold mortar with 8 ml of sodium phosphate buffer (0.02 M – pH 7.4). After the addition of 0.8 g of carborundum, the inoculum sap was rubbed onto the youngest leaves of thirty-day old broad bean plants. The inoculated leaves were rinsed with water 15 min after their inoculation.

### ALCV transmission by aphids

Virus acquisition feedings were performed on broad bean plants 4 to 6 weeks after their agroinfection with ALCV, in a P2 containment chamber under 16h light at 24°C, and 8h dark at 22°C. The duration of the acquisition access period (AAP) was specific to each test (see below the detailed description of experiments) and carried out with 50 individuals per source plants with 1-3 day-old apterous adults. Virus inoculation feedings were performed on 8-day-old broad bean plants under 16h light at 26°C, and 8h dark at 24°C. The duration of the inoculation access period (IAP) and the number of insects transferred to each test plant were specific to each test (see detailed descriptions below). The IAP was stopped by spraying the test plants with Pirimor G insecticide (1 g.l^-1^ in water). The transmission success was assessed by symptom observation and detection of ALCV DNA by PCR as described above.

### Aphid dissection, DNA extraction, and qPCR

After collection, aphids were stored at -20°C until use. Some aphids were dissected to assess the viral content in guts heads and hemolymphs. Dissection and DNA extraction was performed as described by Ryckebusch *et al.* (9).

The qPCR amplification was performed with the LightCycler FastStart DNA Master Plus SYBR Green I kit (Roche) and the LightCycler 480 thermocycler (Roche). Primers ALCV2cEcmLV-F and ALCV2cEcmLV-R (see above) were used at a concentration of 0.6 µM each in a total reaction volume of 10 µl containing 8 µl Master-mix and 2 µl of DNA extract. The cycling protocol and DNA accumulation reports were as described by Ryckebusch *et al.*(9).

### Plant material used for fluorescent in situ hybridization (FISH)

Broad bean and *N. benthamiana* plants were inoculated with agrobacteria containing either an ALCV genome or an empty plasmid as a negative control. ALCV-infected plants were identified four to six weeks after agroinoculation by symptoms and an ALCV-specific PCR test (see above). FISH was performed on petioles or leaves at six weeks after agroinoculation. One cm cross-sections of petioles were cut with a razor blade from upper leaves of ALCV agroinfected broad bean plants. Sections were fixed overnight at 4°C under stirring in embryo dishes containing 4% paraformaldehyde (PFA) diluted in phosphate-buffered saline (PBS). Fixation was stopped by a 15-min incubation in PBS containing 0.1 M glycine. Petiole sections were embedded vertically in 8% low melting agarose in a 24-well tissue culture plate and stored overnight at 4°C. The agarose blocks were extracted from the plates and cross-sections of 100 µm were produced with a Vibratome HM650V (Microm) and processed as described in Sicard *et al.*, (25). Leaf discs of *N. benthamiana* plants were cut with drinking straws and leaf cuticle was removed with forceps. Discs were fixed as described above with petioles. After rinsing in 70% EtOH discs were incubated for 2 hours in Carnoy solution consisting of six volumes of chloroform, three volumes of ethanol, and one volume of acetic acid. Discs were then bleached for 10 minutes in a 6% H_2_O_2_ solution and finally incubated for one hour in PBS before FISH. Vein networks of broad bean leaves were pulled using sticky tapes (26), and subsequently processed like leaf discs.

### Preparation of fluorescent probes and labeling procedure

A fluorescent probe complementary to the CP gene of ALCV was prepared by random priming with the BioPrime DNA labeling system (Invitrogen) and Alexa Fluor 568-labeled dUTP. The template DNA was PCR amplified from the recombinant plasmid containing the ALCV genome, using the following primer pair: CP_ALCV_620-F, 5’-GAA GAG GGC GAG AAC GAC AG-3’ and CP_ALCV_1025-R, 5’-GTG GTC TAT TTC AGC AGT TGC C -3’.

Ten µl of the probe was diluted in 290 µl of 20 mM Tris-HCl hybridization buffer (pH 8) containing 0.9 M NaCl, 0.01% SDS and 30% formamide. The diluted probe was denatured 10 min at 100°C and rapidly cooled on ice for 15 min. In parallel, plant samples (petiole, leaf-discs or veins) were soaked 3 times 5 min in hybridization buffer. Plant samples were then incubated overnight at 37°C in embryo dishes containing probe solutions and sealed with parafilm membranes. After three washing steps of 5 min with hybridization buffer and two with PBS, samples were mounted on microscope slides in Vectashield antifade mounting medium containing 1.5 µg.ml^-1^ DAPI for staining nuclei. Observations were performed using a Zeiss Confocal microscope and acquired in a stack mode.

### EPG system

We used the electropenetrography (EPG) technique (27, 28) to investigate which specific stylet penetration activities of *A. craccivora* individuals were associated with the transmission of the virus. By connecting an insect and its host plant to an electrical circuit, EPG allows variations in biopotentials and electrical resistance to be recorded (referred to as electrical waveforms) and related to different feeding activity patterns (29, 30). In particular, localization of the aphid stylets in the phloem is characterized by two typical waveforms, E1 and E2. The E1 waveform occurring few seconds after the stylet penetration in the sieve element has been related to salivation and described as the essential phase for inoculation of persistently or semi-persistently transmitted plant viruses by aphids (31), whiteflies (32), or leafhoppers (33). The E1 waveform is generally - but not necessarily - followed by a short transition period and typical E2 waveforms. The latter has been related to passive sap ingestion and concurrent secretion of watery saliva (31). An assumption that is commonly accepted is that phloem-restricted viruses transmitted by homopterans are acquired only during the ingestion phase (i.e. waveform E2). Long periods of sustained phloem ingestion would result in a significant increase in acquisition efficiency (34).

A Giga-8 DC-EPG device (EPG-Systems, Wageningen, The Netherlands) was used to monitor probing and ingestion activities of virus-free newly emerged *A. craccivora* adults on leaves of virus-infected broad bean plants. The electrical circuits containing insects, plants, and electrodes were placed in a Faraday cage to isolate them from electromagnetic interference. The electrical signals between the electrodes were converted into digital signals via the Di710-UL (DATAQ, Akron, OH, USA) analog-to-digital board. The digital signals were visualized and recorded on a computer using Probe 3.5 software (EPG Systems, The Netherlands). Recordings were made under constant temperature (23±1°C).

A gold wire (Ø 18.5 µm, 2-3 cm long) was fixed to the thorax cuticle of the backside of the insects using a drop of silver glue (EPG Systems, The Netherlands). During this procedure, insects were held stationary at the tip of a plastic pipe in which a slight suction was applied. The other end of the gold wire was pasted to a copper electrode with the silver glue. This electrode (5 cm long, 2 mm in diameter) was rammed into the soil beside the plant whereas the insect was placed onto a leaf. Before each beginning of recordings, individuals were given a fasting period of 30 minutes. Eight individuals were recorded in parallel. The voltage source was tuned as described by Tjallingii (35), so that the amplifier output signal was between +5 and -5 V, with positive values when stylet tips were outside cells and negative inside cells. After each recording, plants were replaced by new ones.

Signals were analyzed using the software Stylet+a (EPG Systems, The Netherlands). Individuals of many aphid species have been monitored by EPG over the past 50 years, including *A. craccivora* (e.g. 36, 37), allowing an unambiguous interpretation of our own recordings, in particular the waveforms correlated with the stylet location in the phloem (E1 and E2).

### Detection by EPG of vector feeding behavior associated with ALCV acquisition (experiments 1 & 2)

Young apterous adult aphids (⋍2 days old) were given access individually to upper leaves of ALCV-infected broad bean plants and their feeding behavior was monitored by EPG-recording.

In experiment 1, 24 individuals were stopped at the end of their first phloem salivation phase (E1) (modality 1), whereas 30 individuals were monitored up to 4 hours, a period that was expected to contain almost one phloem-feeding phase (E2) (modality 2). Indeed, preliminary analysis revealed that most aphids reached phloem in less than one hour. After the monitoring, aphids were stored individually at -20°C. Total DNA was extracted from each aphid and viral DNA was quantified by qPCR as described above. Negative controls consisted of three adult aphids of the same batch that were given access to upper leaves of healthy broad bean plants.

In experiment 2, 26 individuals were monitored for 4 hours. For each individual, we calculated the variable WDi - Waveform Duration by Insect, i.e. sum of durations of all its events of one waveform type made by each individual insect that produced that waveform - related to E2 pattern as defined by Backus *et al.* (38). The total DNA of each aphid was extracted and viral DNA was quantified by qPCR. Finally, the viral amount of each individual was plotted against WDi-E2.

### Individual transmission probability and transmission rates in relation to aphid numbers (experiments 3 & 4)

In experiment 3, the individual transmission probability of ALCV by aphids was estimated in relation to their virus content. After a 3-day AAP on ALCV-infected broad bean plants, 43 individuals were each given individual access to one healthy broad bean plants for a 5-day IAP. The viral DNA of each of the 34 individuals that were alive at the end of the IAP was quantified by qPCR. The 34 plants associated with the surviving aphids were sampled 4 weeks after IAP and PCR-tested for ALCV detection.

In experiment 4, the transmission rate of ALCV was determined with aphid batches of various sizes in eight independent transmission tests (Suppl Table 1). The batch size were of 1, 5, 10, 20, 30, 40, or 100 individuals depending on the test. The average transmission rate determined for each batch size was used to plot a curve showing the expected transmission rate as a function of the number of individuals per test plants. The theoretical transmission rate *TR* is defined as *TR* = 1-(1-p_i_)^n^ in which p_i_ is the probability that at least one aphid is infective in a population, 1-p_i_ the probability for an aphid not to be infective, and n the batch size.

### Kinetics of ALCV accumulation in aphids (experiment 5)

Young apterous adult aphids were allowed to feed on ALCV infected broad bean plants for 2h, 6h, 15h 24h, 48h and 82h. Individuals were collected at each time point. Their total DNA was extracted in pools of five individuals, and the amount of ALCV DNA was quantified by qPCR. The same test was performed with AAPs of 2h, 4h, 6h, 19h, 24h, 48h, 72h and 96h except that DNA was extracted from individual aphids. Some of the individuals that were not shifted to infected plants for an AAP were collected and tested as negative controls (time point 0h).

### Transmission rates in relation to AAP durations (experiment 6)

Eight independent transmission tests were carried out with 5 or 10 apterous adult aphids per test plant. The transmission rate was determined with 19 or 20 test plants per AAP duration. AAP duration ranged between 1 and 120h and IAP duration was of 5 days.

### ALCV persistence in aphids (experiment 7)

After a 3-day AAP on ALCV infected broad bean plants (D0), apterous adult individuals were shifted to non-infected broad bean plants by groups of 10 individuals per plant. After 4 days, individuals were similarly shifted for two more sequential 4-day feedings on healthy broad bean plants. At D0 and at the end of each sequential 4-day post-AAP feeding (i.e, at D4, D8, D12), 8 groups of 10 individuals were collected. Total viral DNA was extracted from 4 groups per time point and viral DNA was quantified by qPCR. Individuals of the 4 other groups were dissected and the viral DNA was quantified from each pool of 10 organs and 10 hemolymph samples. For each modality, we obtained four independent values of the amount of viral DNA contained in 10 aphids, organs or hemolymph samples. These values were then divided by ten to estimate the mean copy number of viral DNA per insect. For the whole body, the amount of DNA was not measured at D12 because of a limited number of individuals still alive. As a control, individuals from the same rearing as those used for acquisition were analyzed.

### Minimum latency and inoculation period (experiment 8)

In the first set of four transmission tests (tests 1-4), five to ten young apterous adult aphids were given a 15-hours AAP on ALCV infected broad bean plants and then shifted to test plants for various IAP durations between 3h and 48h. Similarly, five to ten aphids were given AAPs of 48 or 72 hours and then shifted to test plants for IAP of various durations between 1 and 120h (tests 5-7). The number of plants for each condition of each test was between 9 and 20. The infectious success was determined with symptom observation and PCR detection of viral DNA.

### Vertical transmission of ALCV (experiment 9)

In a first test, aphids were reared on ALCV infected broad bean plants for 2 weeks. Apterous adult individuals were shifted into a transparent box for 6 hours. Progenies produced in the box were given access to nine healthy plants (5 nymphs per plant). As a positive control of ALCV transmission, adults from the box were given access to three healthy plants (5 apterous adults per plant). The test was repeated independently one month later with 8 healthy plants, each exposed to 5 nymphs. Another positive control was to test if nymphs can transmit ALCV horizontally. To do this, L1-L2 nymphs produced on ALCV infected plants were shifted to broad bean test plants, 5 individuals per plant. Eight day-old adults were similarly shifted to test plants as a control.

Vertical transmission was also evaluated by testing the presence of viral DNA in two pools of 30 nymphs produced by viruliferous adults. Pools were from broad bean plants of the third sequential passage in the persistence test (experiment 7). Total DNA was extracted from each of the two pools of nymphs and the presence of viral DNA was determined by qPCR.

### Effect of ALCV on aphid fitness (experiment 10)

To determine whether ALCV could affect the fitness of its vector, we estimated the intrinsic rate of increase (*r*_*m*_), a parameter combining fecundity and developmental time (39). It was defined as *r*_*m*_ = 0.738.Log_e_(*Md*)/*d*, according to the simplified method of Wyatt & White (40) where *d* is the mean number of days from aphid birth to reproduction (i.e. pre-reproductive time), and *Md* the average number of progeny produced in a time equal to *d.* We also estimated: (i) the mean generation time (*T*), i.e. the mean length of aphid generation, calculated after the approximation of Wyatt & White (40) (*T*=*d*/0.738); and (ii) the doubling time (*DT*), i.e. the time required by the aphid population to double its size, which can be derived from the standard definition of *r*_*m*_ as *DT* = Log_e_(2)*/r*_*m*_ (e.g. (41). The experiment has been conducted with two cohorts of 44 individuals prepared as follows: on day 0 (D0), 2-3-day-old young apterous females from a synchronized population were placed individually on healthy or infected broad bean seedlings for a four-hour laying, and removed thereafter. At day 4 (D4), nymphs were removed except one. From day 6 (D6), individuals were daily observed to detect the age at which each individual lay its first nymphs (i.e. to determine *d*, the pre-reproductive time). Then, the number of nymphs produced during a period of *d* days was counted for each individual. When nymphs retained for laying turned out to be alate, they were excluded from the test. Likewise, the countings that were performed with adults that were not found at the end of the experiment, or that produced very few or no nymphs were excluded from the statistical analysis to avoid interpretation bias. Due to this selection, 6 viruliferous and 9 non-viruliferous individuals were excluded from the analysis. The *r*_*m*_ value was estimated for each individual for comparison of mean differences, but the *r*_*m*_ values presented in the results are the group values, and standard errors were calculated using the bootstrap technique (42).

### Statistical analysis

All statistical analyses were conducted with the R software v3.6.1 (43).

To interpret the results from experiments 2, 5 and 6, we applied *loess* (acronym for locally weighted regression) smoothing to fit a curve through points in each scatterplot (44). With this non-parametric regression technique, no assumptions have to be made about the underlying distribution of the data (45). Smoothed curves were obtained using the *loess* method implemented in the *ggplot2* package of R [function *geom_point()*], with the default values for α=0.75 (this parameter determines the degree of smoothing, i.e. the proportion of all data that is to be used in each local fit), and for λ=2 (i.e. the local regression fitting based on quadratic equations). In experiment 2, Kendall’s tau coefficient was used to measure the ordinal association between cumulative time in E2 and viral load. The test was performed with *cor.test()* function, method *kendall*. To interpret the results from experiments 7, box-plots were used (function *boxplot()* of the package *stats*), and the means were compared using the Kruskal– Wallis rank-sum test (function *krustal.test* of the package *stats*). When the null hypothesis of mean equality was rejected at the 5% threshold, the means of each pair of modalities were compared using the multiple comparison method based on the Benjamini and Yekutieli (46) procedure (function *pairwise.t.test()* with p-value adjustment method: BY). The means of the parameters estimated in experiment 10 were compared using the Wilcoxon rank-sum test (function *wilcox.test()* of the package *stats*). Standard errors of these parameters were calculated using a bootstrap procedure with the *boot()* function of the *boot* package.

## RESULTS

### ALCV is phloem restricted and heterogeneously distributed in this tissue compartment

A fluorescent probe complementary to the CP gene of ALCV was used to localize ALCV in broad bean plants by FISH (Fig. 1). Fluorescent labeling was detected in petioles of plants agroinfected with ALCV (Fig. 1a) but not in petioles of mock-inoculated plants. Specific labelling co-localized with cell nuclei and was detected in relatively few cells, with a maximum of 3 labeled cells per cross-section. Specific labelling was restricted to areas located between xylem and sclerenchyma, the location of phloem (Fig. 1a,b). ALCV specific labelling was detected also in phloem of vascular tissues pulled from broad bean leaf lamina (Fig. 1c). The labelling is restricted to some sections of the vascular network and only some cells of these sections are labelled. The specifically labelled cells are nucleated and characterized by an elongated shape. They may be identified as phloem parenchyma cells or companion cells. Translaminar observations of ALCV-infected *N. benthamiana* leaves showed FISH labelling in phloem but not outside the vascular network (Fig. 1f). Consistently with these results suggesting that non-phloem cells are immune to ALCV, none of the mechanically inoculated plants (0/37) displayed ALCV symptoms or were detected PCR-positive for the presence of ALCV.

**Figure 1.**
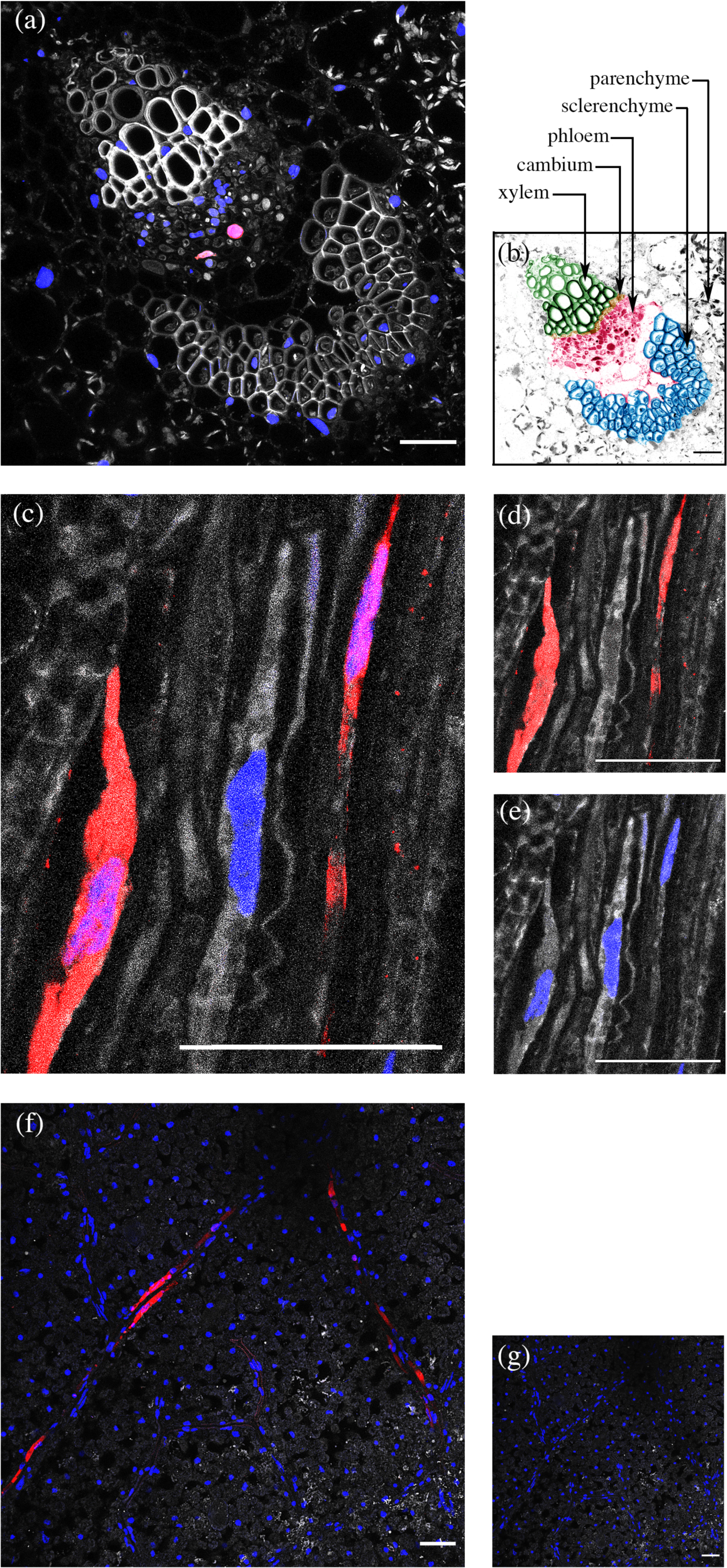
Histological localization of ALCV DNA in plants tissues by FISH. The ALCV specific DNA probe is labeled with red Alexa (568) fluorochrome. Nuclei are DAPI-blue stained. Color channels were merged in panels a, c, f and g, but not in panels d and e. (a) Cross section of a petiole of an ALCV-infected broad bean plant, showing a vascular bundle (b). Same as (a) but with artificial colors: xylem in green, cambium in yellow, phloem in pink, sclerenchyma in blue, parenchyma in grey. (c, d, e) Elongated cells of vascular bundles pulled from a leaf of an ALCV-infected faba bean plant. (d) Same as (c) but only with the red channel. (e) same as (c) but only with the blue channel. (f) Translaminar view of a leaf sampled on an ALCV-infected *Nicotiana benthamiana* plant. (g) Same as (f) except that the leaf was collected on a healthy plant. Preparations were examined with confocal microscopy. Horizontal bars = 50 µm.

ALCV distribution was also investigated by the characterization of plant tissues where aphids can acquire the virus (experiment 1). To do this, we monitored the feeding behavior of 54 aphid individuals on ALCV-infected plants by EPG and tested whether particular electrical signals may be associated with virus acquisition. ALCV was not qPCR detected in the 24 individuals for which the EPG records were stopped at the end of their first phloem salivation phase, i.e E1. This particular waveform is recorded when individuals introduce their stylets in sieve tubes but without sap ingestion (Fig. 2c). ALCV was neither detected in the 10 individuals for which no phloem waveforms were observed (E1 or E2), which indicates that only non-phloem cells were probed. On the contrary, among the 20 individuals that reached E2, the waveform associated with phloemian sap ingestion, 14 were detected ALCV-positive by qPCR. Of the six insects detected negative for ALCV detection, five remained less than 20 min in phase E2, a duration that may have been too short to acquire a detectable amount of viral DNA. Nevertheless, one aphid has been in the phase of phloem sap ingestion for more than 2h30 and has not acquired the virus either. FISH and EPG results are consistent with each other because both show that ALCV is phloem restricted and that ALCV is not uniformly distributed in this tissue.

**Figure 2.**
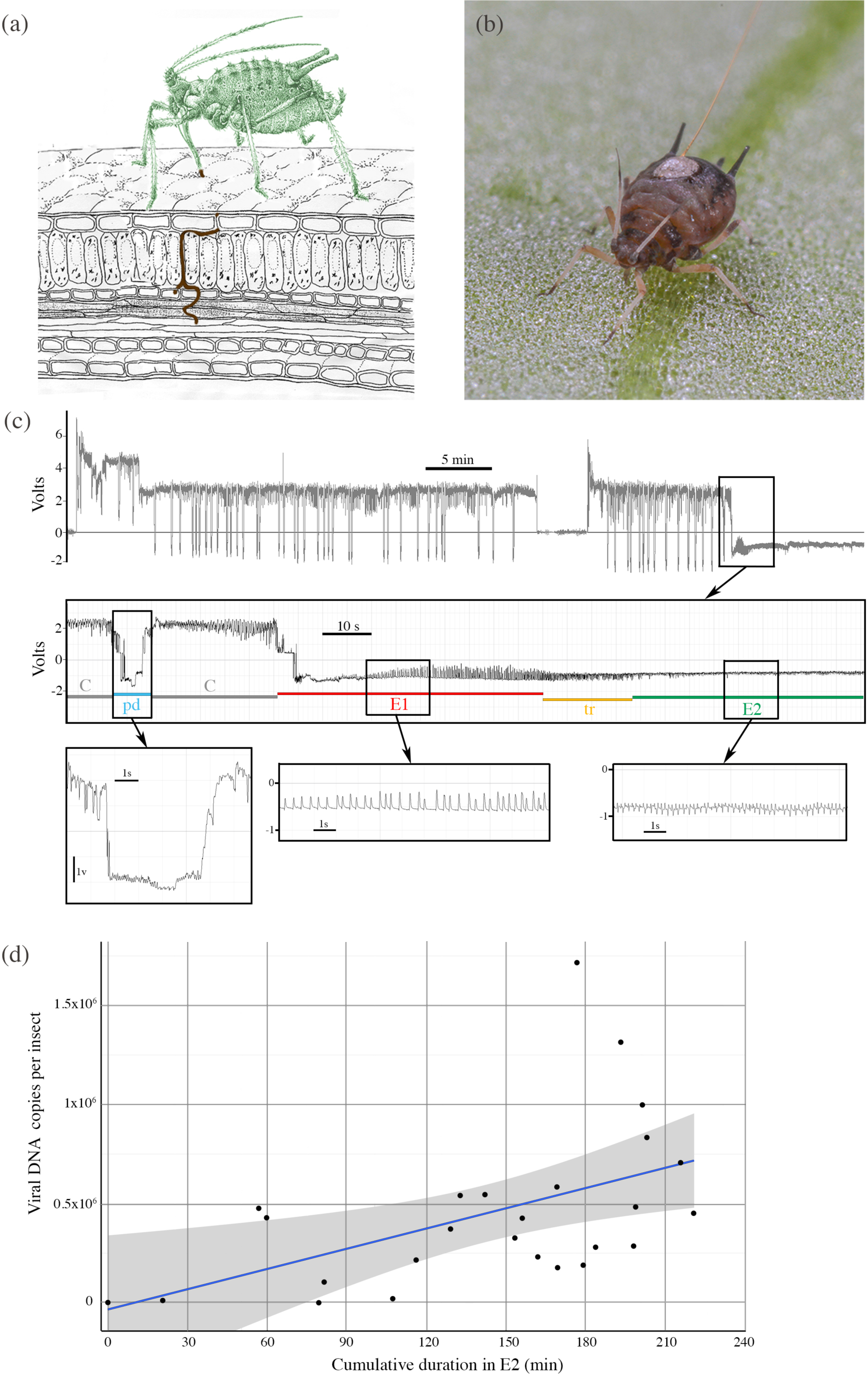
EPG design, and *Aphis craccivora* feeding behavior associated with ALCV acquisition. (a) Schematic illustration of an aphid in feeding position showing the intra-leaf route of its stylets until a vascular bundle; (b) *A. craccivora* adult with a thin gold wire glued to its dorsum with a small drop of silver print paint; (c) Overview of typical EPG waveforms produced by an apterous adult feeding for 1h on a branch of broad bean plant, and expanded views of the waveforms studied: pd (intracellular puncture in epidermis or mesophyll cell), E1 phase (salivation into sieve tube elements of the phloem), E2 phase (ingestion from sieve tube elements), and tr (transition between E1/E2); (d) Cumulative duration in phase E2 of young apterous adults measured during 4 h EPG recordings on ALCV infected branches of broad bean plants, plotted against their post-EPG viral DNA content (experiment 2). Smoothing method was used to add a regression line with 0.95 confidence interval shown in grey. (black dots inside the envelope, red dots outside the envelope).

### Efficiency of acquisition is dependant on the puncture site and AAP duration

The heterogeneous phloem distribution of ALCV revealed by FISH and EPG suggests that the efficiency of virus acquisition by aphids is dependant on the puncture site. This hypothesis was validated with a set of adult aphids for which we accurately assessed both the duration of phloem ingestion by EPG (E2 waveform) and the acquired viral DNA content by qPCR (Fig. 2d). Indeed, there is no correlation between E2 cumulative time and the amount of virus ingested by the aphid (Kendall’s rank correlation thau, τ = 0.0027). For example, two aphids for which the E2 waveform was recorded for about one hour, acquired each as many or more viral DNA copies (0.5×10^6^) than aphids for which E2 recording lasted more than 2h30. On the other hand, one aphid for which E2 recording lasted almost two hours in phase E2 has acquired very little viral DNA. Nevertheless, it is noteworthy that individuals that fed more than 107 min were all detected qPCR positive with more than 10^5^ copies of viral DNA, suggesting that in spite of heterogeneous distribution, the virus is apparently accessible at any sites of the phloem network.

This result suggests that phloem-feeding durations and viral DNA accumulation may be positively correlated with longer AAP durations. This prediction was validated by monitoring viral DNA accumulation in individuals that were given access to ALCV infected plants for durations ranging between 2 and 96 hours (Fig. 3). Aphids accumulate a large amount of virus in the first 15-19 hours of acquisition. During this initial period, viral load per insect increased sharply up to 4-7×10^5^ copies. Thereafter, the viral increase was lower with only 10 times increase up to a plateau of about 2 to 7×10^6^ copies that were reached at 48 hours. Interestingly, aphids individuals that were given a 2-hour AAP were not all PCR-positive in spite of the fact that they were tested in pools of 10 individuals (4 of the 12 pools that were given a 2-hour AAP were negative). This result is consistent with the heterogeneous virus acquisition observed with the EPG-monitored individuals that were given 4 hour AAPs (Fig. 2d).

**Figure 3.**
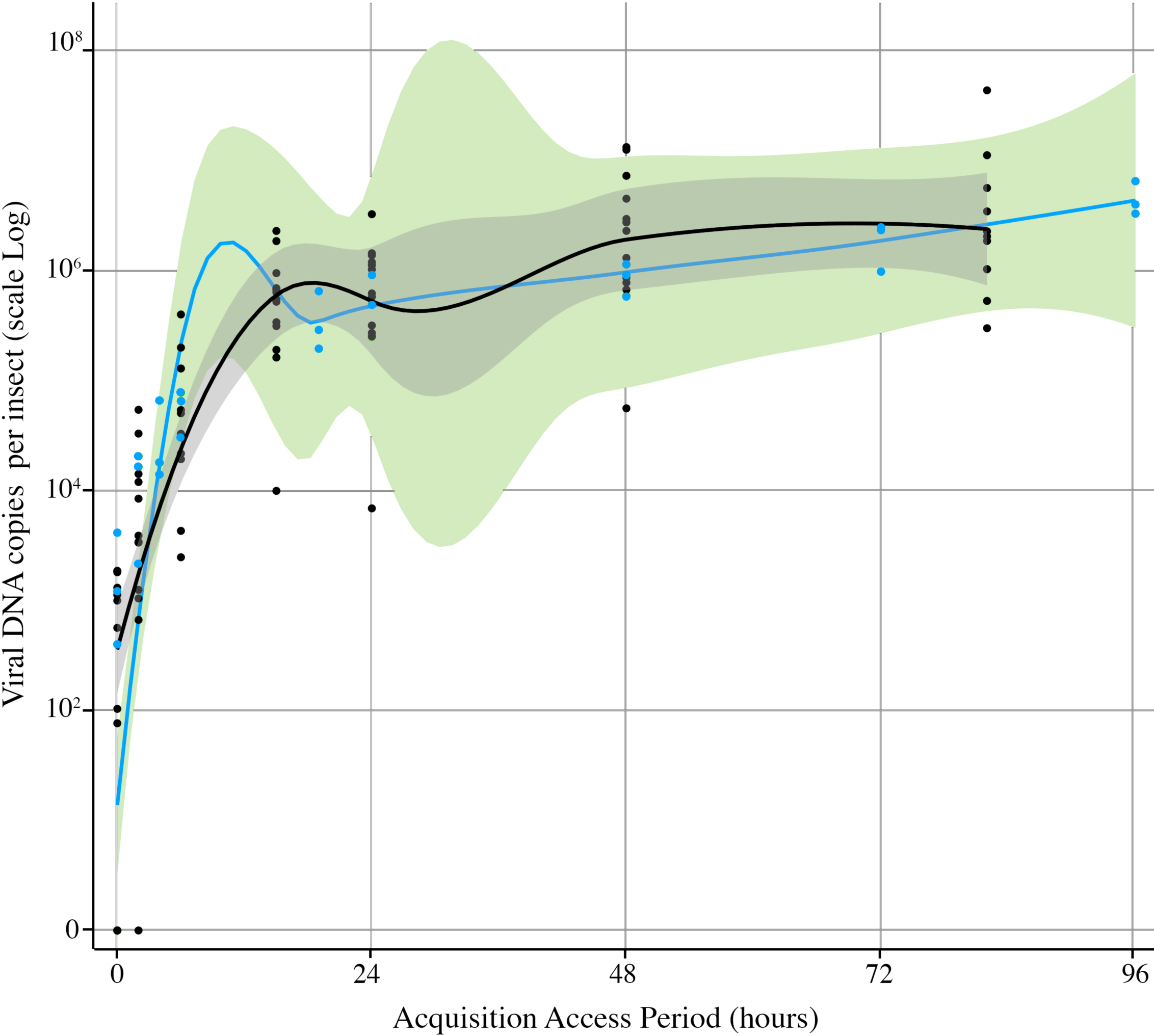
Kinetics of ALCV accumulation in *Aphis craccivora*. Viral accumulation was assessed by monitoring viral DNA contents with qPCR on individuals that were given AAPs of 2 to 82 h (black dots) or 2 to 96 h (blue dots). While black dots correspond to individuals analysed individually, each blue dot corresponds to the average viral content of 5 individuals analysed by qPCR in a pool. For each set of dots, a smooth local regression was performed, in black for the black dots and in blue for the blue dots. Moreover a 0.95 confidence interval was displayed around smooth in grey and green colors respectively.

### A high accumulation of ALCV is necessary for efficient transmission but not always sufficient

The accumulation dynamics of ALCV in aphids revealed that viral DNA content reaches more than 10^6^ viral DNA copies per insect within two days of AAP (Fig. 3). However, it was not known if this acquired virus can be easily transmitted and particularly if the transmission is subjected to an accumulation threshold. To test the threshold hypothesis, 43 individuals, following a 3-day AAP, were given access to test plants - one test plant per individual - to determine their ability to transmit the acquired virus (Fig. 4a). The viral amounts assessed by qPCR in the 34 individuals collected alive at the end of the 5-day IAP were consistent with the threshold hypothesis. Indeed the four individuals that were able to transmit ALCV to their test plant were among the eight individuals that exhibited the highest virus content, i.e above 1.6 × 10^7^ viral DNA copies. Aphids whose viral amount was below this threshold failed to transmit ALCV. The 4 non-transmitters whose viral content was above the threshold suggest that, although necessary, accumulating virus above a threshold is not always sufficient to transmission. Other parameters, presently unknown, might also determine the efficiency of viral inoculation.

**Figure 4.**
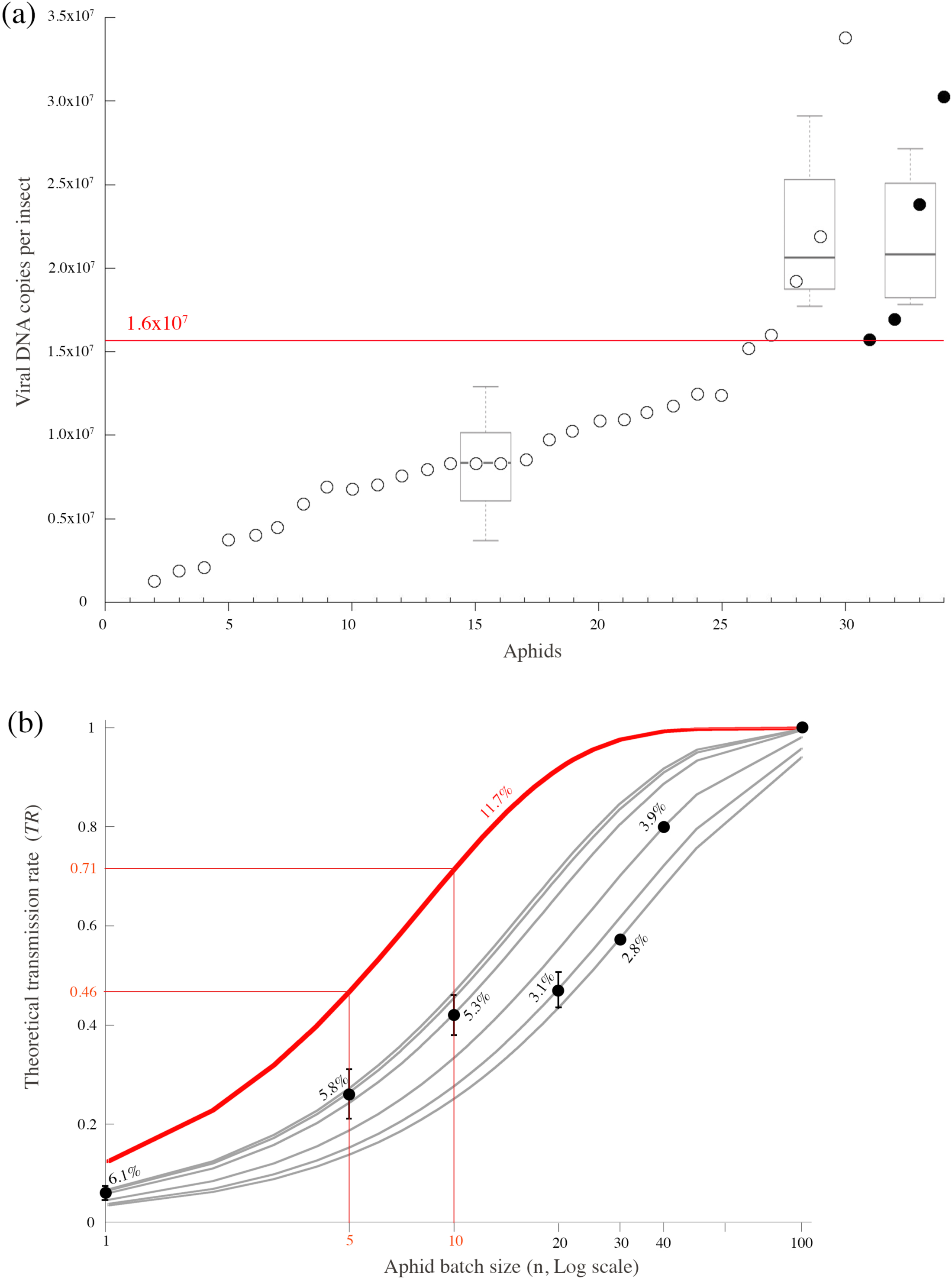
Transmission rate of ALCV by *A. craccivora* individuals and relationship with their viral amount (a) and the number of individuals per test plant (b). (a) Viral DNA amount assessed by qPCR in 34 individuals after a 3-day AAP on ALCV-infected broad bean plants and a 5-day IAP on healthy plants with one individual per test plant. While solid circles represent individuals that transmitted ALCV to their test plant, open circles represent non-transmitters. Only individuals in which viral content was above 1.6 × 10^7^ viral DNA copies (red horizontal line) transmitted ALCV. (b) Transmission rates were determined experimentally (black dots with standard deviations) in seven independent transmission tests performed with 1, 5, 10, 20, 30, 40, 100 individuals (experiment 4). For each aphid batch size, mean theoretical individual transmission rate (pi) was deduced from formula, *TR* = 1-(1-pi)^n^, knowing *TR* and n (values in % on the figure) (see also Suppl Table 1). The grey curves are the theoretical curves of *TR* as a function of n for these pi values. The red curve was derived from the transmission test shown in (a) in which p_i_=4/34.

The individual transmission rate (p_i_⋍11.8%; 4/34) is relatively low considering the long durations of AAP and IAP, - 3 and 5 days respectively. Using p_i_⋍11.8%, theoretical transmission rates (*TR*) were estimated as a function of the number of individuals used per test plant according to the formula *TR* = 1-(1-p_i_)^n^ (Fig 4b, red curve). Surprisingly, the transmission rates derived from transmission tests performed with different numbers of individuals per test plants (1, 5, 10, 20, 30, 40; Suppl Table 1), were all below the expected rates (Fig. 4b). The theoretical individual transmission rates derived from the observed transmission rates with the formula *TR* = 1-(1-p_i_)^n^ were between 2.8 and 6.1%. To summarize, the individual transmission rate of ALCV by *A. craccivora* is generally around 4-5% with a maximum of 12% when the transmission was performed with one insect per test plant. Although the increase of transmission rate is positively correlated with the increase of viruliferous individuals, the transmission success cannot be easily predicted from individual transmission rates. Increasing the number of aphids per test plant may influence the complex plant-virus-insect interactions, for example by increasing plant defenses, and in turn, reduce transmission efficiency. In view of these results, a convenient compromise to reach an amenable transmission rate (18-45%) with a minimum number of insects and test plants, is with 5 or 10 individuals per test plant.

### Optimal transmission rate needs a minimum AAP duration of 48 hours

To determine the minimum AAP duration that is needed to reach an optimal transmission rate, 5 or 10 individuals per test plants were used as defined above. Figure 5 shows that up to 48h, the transmission rate is a quasi-linear function of acquisition time, reaching a maximum of about 50 to 60%. Increasing acquisition time beyond 48h does not increase the transmission rate, even up to 5 days. It is noteworthy that the maximum transmission rate is not reached following a 24 hour AAP although the most intensive virus accumulation was achieved in less than 24h (Fig. 3). These results are consistent with the threshold hypothesis. Indeed, it seems that the moderate virus accumulation that occurs after the initial intensive virus accumulation (19h) is critical for an optimal transmission rate. Unexpectedly, in some tests, low transmission rates were observed although the experimental conditions were the same.

**Figure 5.**
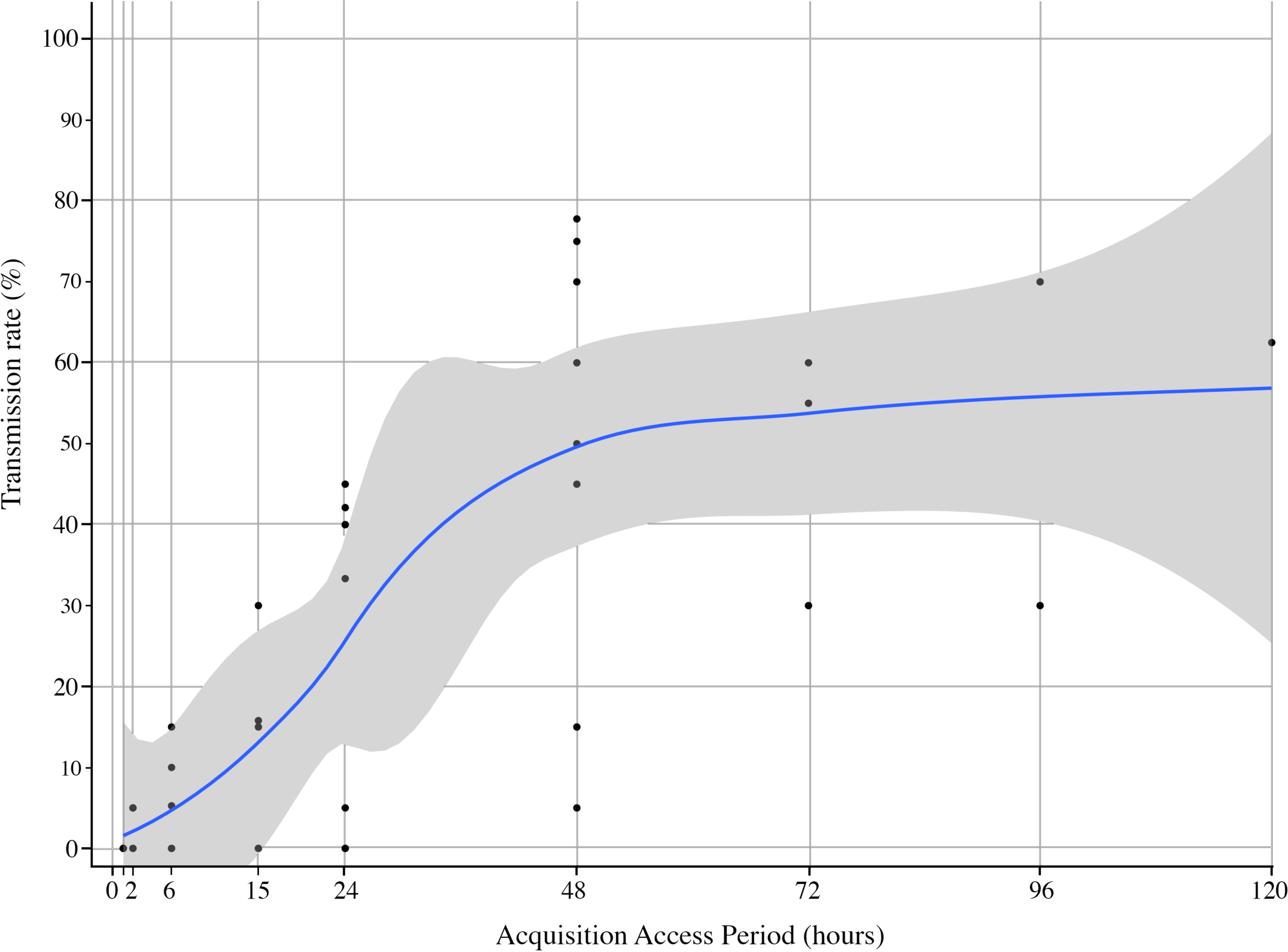
Relationship between AAP duration and transmission rate of ALCV by *Aphis craccivora*. Each dot corresponds to a transmission rate determined with 5 days AAP carried out with 19 or 20 test plants and with 5 or 10 individuals per test plant. The figure summarizes the results generated with eight independent transmission tests (experiment 6). A smooth local regression was performed and 0.95 confidence interval was displayed around smooth.

### Contrasted ALCV persistence in aphid cellular compartments, and potential impact on virus inoculation

ALCV was previously reported to circulate and persist in *A. craccivora* based on the detection of ALCV DNA in midguts, heads and hemolymphs 6 days after a 3 day AAP (9). This result is confirmed and expanded here by monitoring the dynamics of viral DNA in these compartments up to 12 days after a 3 day AAP (Fig. 6). Viral DNA monitored from the whole body exhibited an increase between 0 and 8 days post-AAP and the effect of time was significant according to the Kruskal-Wallis test (p-value=0.048). However, the pairwise multiple comparison test does not reject the null hypothesis of equality of means estimated for every three post-AAP durations (p-values ≥ 0.14). This apparent contradiction between the tests can be explained by the low number of repetitions (n=4) for each duration. We conservatively conclude that the viral load in the whole body did not increase after AAP which is consistent with the midgut results in which no significant differences were detected between the samples collected over time (Kruskal-Wallis test, p-value=0.653). In hemolymph and heads, the viral amount exhibited a decrease over time (Kruskal-Wallis test, respectively p-value=0.047 and p-value=0.034). The pairwise multiple comparison test further support this decrease in hemolymphs. Indeed, a significant difference in viral content was detected between 0 and 12 days post-AAP (p-value=0.016). One of the 4 hemolymph samples collected at 12 days post-AAP was negative for viral DNA detection. The pairwise test did not show any significant difference in viral content in the heads (p-value=0.098). Nevertheless, one of the 4 head samples was negative at 9 and 12 days post AAP.

**Figure 6.**
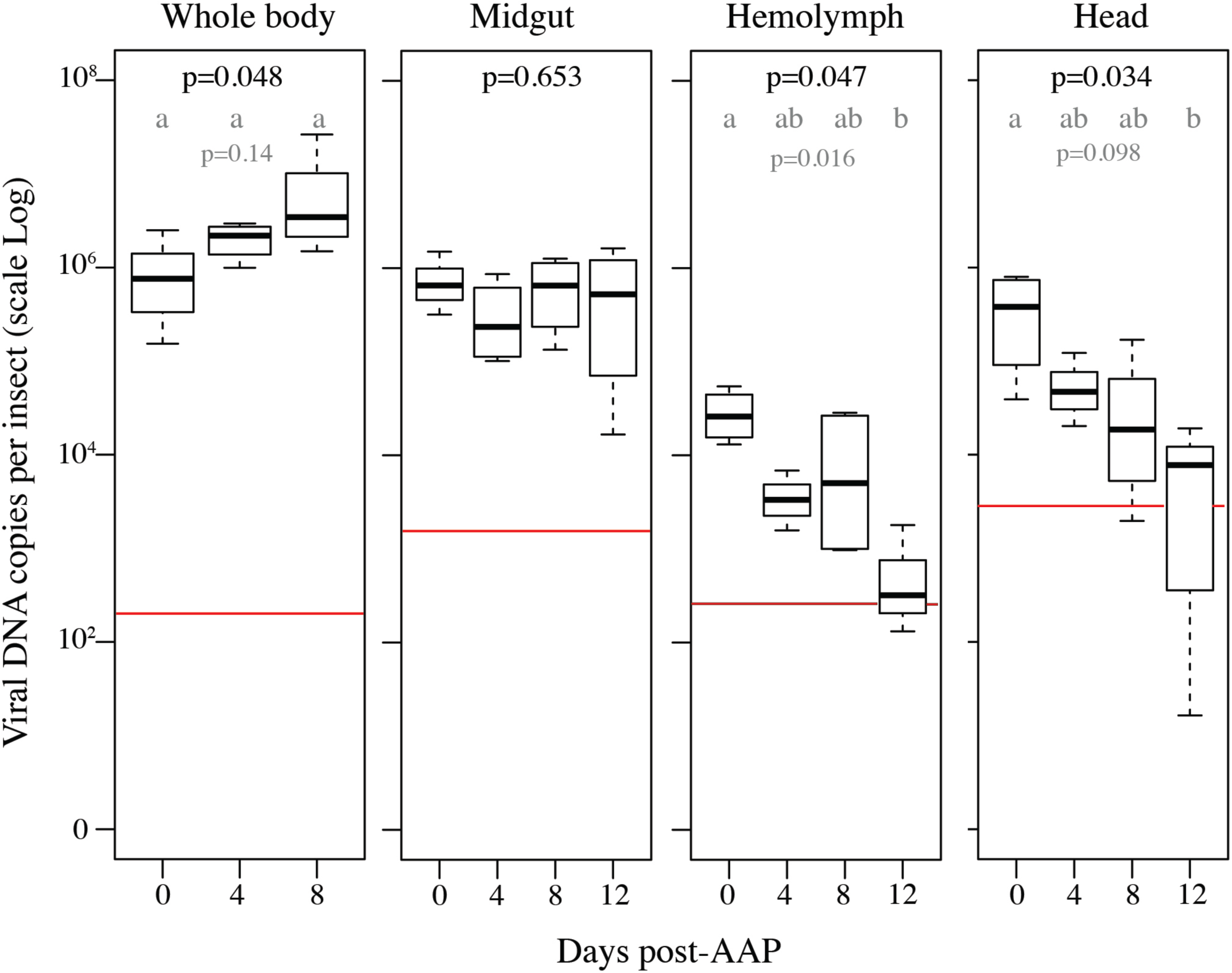
ALCV persistence in *Aphis craccivora* cellular compartments. The box-plots show the amount of ALCV DNA in whole body, midgut, hemolymph, and head of aphids following a 3-day AAP on ALCV infected broad bean plants (D0), and after three sequential 4-day post-AAP feedings (i.e, D4, D8, D12). The content of ALCV DNA was determined by qPCR in 4 pools of 10 non-dissected individuals and 4 pools of 10 organs or hemolymph. Accumulations are reported as logarithm 10 of the number of viral DNA copies. The red lines represent the highest value obtained from individuals sampled before the 3-day AAP. The amount of DNA in non-dissected individuals was not measured at D12 because there were not enough alive aphids on that date. The means for each duration were compared by a Kruskal-Wallis test. When the null hypothesis of mean equality was rejected (p<0.05, p-values in black), the means of each pair of modalities were compared using the multiple comparison method based on Benjamini and Yekutieli’s procedure. Different letters indicate significant differences at the level shown in grey.

The decrease of viral content in the hemolymph and the head compartment was thought to limit the amount of virus that is potentially released from the insect through salivary glands, and hence, the transmission rate. To test the effect of IAP duration on the transmission rate, we used a 48- or 72-hour AAP which, according to previous tests produced viruliferous insects that exhibit optimal infectivity (Fig. 5). With these experimental conditions, it was expected that the virus accumulation would not be a limiting factor for virus inoculation. However, in spite of AAP durations compatible with high virus accumulation, the inoculation was successful only with an IAP duration of at least 24 hours (Table 1). Additional transmission tests were performed to investigate the minimum latency period. To do this we had to use individuals that were given an AAP that was long enough to produce infective aphids (Fig. 5) and short enough to limit the risk of exceeding the potential minimum time of latency during AAP. Using 15 hours AAPs (Table 1), no virus transmission was obtained with IAPs of 3 hours (0 out of 10 test plants), 5 hours (0/40), 6 hours (0/10) and 9 hours (0/40). One of the 4 tests (i.e. test 1) performed with a 12-hour IAP was successful (1/20), which indicates that the minimum latency is of 27 hours or less. Interestingly, this latency test revealed also that the minimum successful IAP may be as short as 12 hours.

**Table 1.** Description of the seven transmission tests designed to estimate the minimum latency and inoculation periods (experiment 8). Ratios between the number of infected plants and the number of test plants are indicated for each test plants and transmission conditions. The infection status of the test plants was determined with symptoms and/or the detection of ALCV DNA by PCR. AAP, acquisition access period; IAP, inoculation access period; n, number of aphid individuals per test plant.

| AAP<br>(h) | IAP<br>(h) | Latency<br>(h) | Test 1<br>n=5 | Test 2<br>n=5 | Test 3<br>n=10 | Test 4<br>n=10 | Test 5<br>n=5 | Test 6<br>n=10 | Test 7<br>n=10 |
| --- | --- | --- | --- | --- | --- | --- | --- | --- | --- |
| 15 | 3 | 18 |  |  |  | 0/10 |  |  |  |
| 15 | 5 | 20 |  | 0/20 | 0/20 |  |  |  |  |
| 15 | 6 | 21 |  |  |  | 0/10 |  |  |  |
| 15 | 9 | 24 |  | 0/20 | 0/20 |  |  |  |  |
| **15** | **12** | **27** | **1/20** | 0/20 | 0/20 | 0/10 |  |  |  |
| **15** | **24** | **39** | **1/20** | 0/20 | 0/20 | 0/10 |  |  |  |
| 15 | 48 | 63 | 0/20 |  |  | 0/12 |  |  |  |
| 48 | 1 | 49 |  |  |  |  |  | 0/10 |  |
| 48 | 6 | 54 |  |  |  |  |  | 0/10 |  |
| 48 | 12 | 60 |  |  |  |  | 0/11 | 0/10 |  |
| **48** | **24** | **72** |  |  |  |  | **2/12** | 0/10 |  |
| **48** | **48** | **96** |  |  |  |  | **4/10** | **1/10** |  |
| 48 | 120 | 168 |  |  |  |  |  | 0/10 |  |
| 72 | 3 | 5 |  |  |  |  |  |  | 0/10 |
| 72 | 6 | 78 |  |  |  |  |  |  | 0/10 |
| 72 | 12 | 84 |  |  |  |  |  |  | 0/10 |
| 72 | 24 | 96 |  |  |  |  |  |  | 0/9 |
| 72 | 48 | 120 |  |  |  |  |  |  | 0/10 |
| **72** | **120** | **192** |  |  |  |  |  |  | **2/10** |

### No vertical transmission

The relatively low efficiency of virus inoculation might be balanced by a potential vertically transmission. To test the hypothesis, nymphs produced by viruliferous aphids were tested either for virus presence or for infectivity on broad bean plants. In the three independent tests, nymphs were not detected PCR-positive for ALCV DNA and did not induce ALCV infection in broad bean plants. The viruliferous parental aphids which were tested as positive controls induced infection in two of the 5 test plants exposed. L1-L2 nymphs produced on ALCV infected plants transmitted ALCV as efficiently as adults from the same plants, with transmission rates of 53% (8/15) and 50% (5/10), respectively.

### No ALCV-associated fitness cost for the host aphids

The fitness of viruliferous and non-viruliferous aphids was compared by estimating the intrinsic rate of natural increase (*r*_*m*_) (Table 2). The mean time of development of non-viruliferous individuals was not significantly different from that of viruliferous individuals (*d*=7.88 days *versus* 7.39 days, Wilcoxon test p-value>0.1). Likewise, the average number of nymphs laid during a period equal to the time of development (*d*) was not significantly different between viruliferous and non-viruliferous individuals with 55.17 and 49.79 nymphs, respectively. Hence, the *r*_*m*_, the combination of both parameters, was highly similar between non-viruliferous and viruliferous individuals (0.418 *versus* 0.422 progenies/female/day).

**Table 2.** Comparing the fitness of *A. craccivora* individuals that acquired ALCV to that of non-infected individuals. *d*: mean number of days from aphid birth to reproduction (i.e. pre-reproductive time); *Md*: average number of progeny produced in a time equal to *d*; # larvae: mean number of nymphs produced per day during a time equal to *d*; *r*_*m*_: intrinsic rate of increase; *T*: mean length of aphid generation; *DT*: time required by the aphid population to double its size; SE: bootstrap estimator of standard error.

| Estimated parameters |  | non-viruliferous aphids<br>N=35 | viruliferous aphids<br>N=38 |
| --- | --- | --- | --- |
| *d* | day (SE) | 7.00 (0.04) | 6.74 (0.08) |
| *Md* | day (SE) | 55.17 (2.63) | 49.79 (2.69) |
| # nymphs | per day (SE) | 7.88 (0.38) | 7.41 (0.40) |
| *r<sub>m</sub>* | prog/fem/day (SE) | 0.418 (0.006) | 0.422 (0.009) |
| *T* | day | 9.49 | 9.13 |
| *DT* | day | 1.64 | 1.62 |

**Table 2.**
| Estimated parameters |  | non-viruliferous aphids | viruliferous aphids |
| --- | --- | --- | --- |
|  |  | N=35 | N=38 |
| *d* | day (SE) | 7.00 (0.04) | 6.74 (0.08) |
| *Md* | day (SE) | 55.17 (2.63) | 49.79 (2.69) |
| # nymphs | per day (SE) | 7.88 (0.38) | 7.41 (0.40) |
| *r<sub>m</sub>* | prog/fem/day (SE) | 0.418 (0.006) | 0.422 (0.009) |
| *T* | day | 9.49 | 9.13 |
| *DT* | day | 1.64 | 1.62 |

## DISCUSSION

Surprisingly aphid-transmitted geminiviruses were detected less than 10 years ago, whilst leafhopper and whitefly transmitted geminiviruses were discovered more than 100 years ago (3, 5, 6, 9, 17). We hypothesized that the delayed discovery of aphid-transmitted geminiviruses is due to their low dissemination rate within cultivated plants compared to that of non-aphid transmitted geminiviruses, which in turn may be associated with potentially low efficiency of transmission by aphids. To test this hypothesis, we determined the transmission parameters of an aphid-transmitted geminivirus and compared them to previously reported transmission parameters of non-aphid transmitted geminiviruses.

### ALCV is restricted and heterogeneously distributed in the phloem tissues

The process in which an insect vector acquires the virus from a plant involves three partners. To monitor these tripartite interactions we used an original approach comprising three complementary techniques. While FISH determined that ALCV was restricted and heterogeneously distributed in the phloem network, the combined EPG and qPCR monitoring were able to accurately assess how efficiently *A. craccivora* individuals acquire ALCV from this plant compartment. Thus, with EPG it was possible firstly to determine that ALCV can be acquired at a detectable level only from the sieve elements of the phloem. Secondly, in association with the accurate quantification of viral DNA in the vector, EPG recordings provided a compelling confirmation of the heterogeneous distribution of ALCV in the phloem network. The fact that all individuals that ingested phloem sap for more than 100 minutes were detected ALCV positive, irrespective of their puncture site, is consistent with generalized contamination of the sieve tubes. The individuals who accumulated the highest concentration of viral DNA are thought to be those for which the punctured sieve element was the closest to virus replicating phloem cells.

Although geminivirus infected plant tissues were frequently tested for their phloem restriction (e.g., 47), heterogeneous virus distribution within the phloem network was to our knowledge never mentioned. However, as most of these microscopic observations were performed on cross-sections, the uneven distribution, if any, was probably not easily detectable. One of the only studies in which longitudinal sections of geminivirus infected plant tissues were performed was with the phloem restricted tomato yellow leaf curl virus (TYLCV), and the reported photos seem to be consistent with heterogeneous distribution (48). This interpretation is indirectly supported by the uneven amount of TYLCV acquired by the whitefly vector *Bemisia tabaci* (49). It is noteworthy that this uneven distribution was detected during a 4-hour AAP, the same duration as in our ALCV assay in which the duration of sieve element access was not correlated to the amount of ALCV in *A. craccivora* individuals. However, as feeding behavior was not monitored in the TYLCV study, it is not known if the heterogeneous TYLCV acquisition is only due to heterogeneous virus distribution or if heterogeneous durations of sieve tube feedings have also contributed to it.

### Heterogeneous intra-phloem distribution of ALCV does not prevent the efficient acquisition

The non-correlation between the amount of acquired ALCV and the durations of AAP observed during a 4-hour AAP fades with longer AAPs. Thus, we hypothesize that, due to the generalized contamination of the sieve tube network, the ALCV content of individual aphids, irrespective of their puncture spot, increase steadily during the first 24 hours of AAP, the accumulation phase exhibiting the highest increase. Such a correlation between duration of AAP and virus amount in the vector was also detected with other geminiviruses and is a common feature of non-propagative transmission (50).

From 15 hours AAP, the number of copies of viral DNA per individual reaches 10^6^, and was in the range of 10^6^ to 10^7^ between 2 and 4 days AAP. While these amounts are slightly higher than that of MSV in its leafhopper vector following a 6-day AAP (< 10^6^ DNA copies)(51), they were lower than those of tomato yellow leaf curl virus and watermelon chlorotic stunt virus in their whitefly vector following AAPs of 5 or 6 days (10^8^ DNA copies or more) (52, 53)

### Transmission success depends on a high viral amount threshold in aphids

Previous transmission studies with geminiviruses showed that the infectivity of viruliferous vectors was correlated with AAP durations (e.g., 54). These results suggest that insect infectivity depends on its virus content but to our knowledge, a threshold of virus concentration beyond which an infection is possible was to our knowledge never defined. Here, by checking the virus concentrations in individual viruliferous aphids each tested for infectivity, we came up with a virus concentration threshold below which insects are not infective. As the quantification was done after the 5-day IAP, the threshold of 1.6 × 10^7^ copies of viral DNA corresponds to the persistent virus that is most likely internalized and therefore relevant for infectivity.

Although necessary for infectivity, reaching this concentration threshold is not sufficient, because 50% of the individuals that reached the threshold were not infective during the 5-day IAP. As aphids are genetically highly homogeneous due to their clonal multiplication, the contrasted infectivities may be associated with stochastic phenomena, like the site of inoculation, a particular feeding behavior, particular physiological conditions influencing the virus distribution along the transmission route. Moreover, the influence of endosymbionts cannot be excluded from individuals that may potentially exhibit contrasting infection status (55).

Interestingly, the infection rates that were obtained with 5 or 10 individuals per test plant following increasing AAP durations can be interpreted according to a critical threshold of ALCV DNA content which is around 10^6^ viral DNA copies per individual (Fig. 3). Indeed, the maximum infection rate, around 50%, was reached only from 48 hours AAP (Fig. 5), which according to the accumulation dynamics of ALCV in *A. craccivora* (Fig. 3) is the time needed for a majority of individuals to reach viral contents that are higher than 10^6^ copies. Thus, although the ALCV content of 24-hour AAP individuals was only slightly lower than that of 48-hour AAP individuals, the differential transmission rate is of about 2. These results suggest that the ALCV content threshold that was found to be necessary for individual aphid transmission cannot be achieved collectively, by the addition of individual contributions. This hypothesis was supported by the low transmission rates obtained with groups of aphids ranging from 5 to 40 individuals. Indeed, they were all below the expected (theoretical) transmission rate calculated from the observed individual transmission rate (12%) (Fig. 4b). Indeed, if this threshold could have been collectively achieved the transmission probability would have been higher than the rate estimated from the probability of having at least one individual per group.

### The low infectivity of aphids is associated with the low persistence of ALCV in hemolymph and head

The highest individual transmission rate of ALCV by its vector *A. craccivora* was 12%, an obviously low rate compared to other circulative transmitted viruses, with for example 90% with the leafhopper transmitted geminivirus MSV (56) and the whitefly transmitted geminivirus TYLCV (57)and 60% with potato leafroll virus (PLRV) (genus *Polerovirus*, family *Geminiviridae*) transmitted by *Myzus persicae* (58). As acquisition dynamics and maximum viral content of ALCV in *A. craccivora* individuals were found to be similar to those of non-aphid transmitted geminiviruses, other parameters must be incriminated to explain the low transmission rate of ALCV. When ALCV DNA was monitored for its retention in dissected individuals or midguts, no decrease was detected up 8 days post AAP. However, ALCV DNA was not efficiently retained in hemolymph and heads which indicate that the low transmission of ALCV is not due to a barrier at the entrance of gut cells and persistence in there, but to incomplete barriers located beyond this stage, including potentially, gut exit, resistance to degradation in hemolymph, salivary gland penetration and salivary gland exit. Interestingly, such barriers to circulative transmission beyond virus persistence in the midgut (post-gut barriers) were detected with other insects/virus combinations associated with low or no transmission. Firstly, post-gut barriers detected with a non-vector population of *A. craccivora* are mostly beyond the stage of virus persistence in the gut (9). Secondly, post-gut barriers were suspected in *Macrosiphum euphorbiae* aphid individuals that transmitted potato leafroll virus (PLRV, genus *Polerovirus*, family *Luteoviridae*) much less efficiently than *Myzus persicae* individuals (59). As the low transmission of PLRV by *M. euphorbiae* could not be ascribed to failure to acquire or retain PLRV, or to degradation of virus particles in the aphid, it was thought that only a few PLRV particles pass from the hemolymph to saliva in this species. Thirdly, post-gut barriers were detected in one of the rare begomovirus/*B. tabaci* combinations in which no transmission was detected, i.e. tomato leaf curl China virus (TYLCCNV) tested with the Mediterranean cryptic species (MED) (60). Although the acquisition and retention of TYLCCNV in the midguts of MED whiteflies were as efficient as that of whiteflies of the MEAM cryptic species (Middle East Asia Minor 1) that are vectors of TYLCCNV, penetration into the primary salivary glands (PSG) was much lower in MED than in MEAM whiteflies. Consistently with this last result, the TYLCCNV was not detected in the saliva of MED individuals. In line with the TYLCCNV results, further studies are needed to test if post-gut barriers to ALCV in *A. craccivora* are also associated with the last transmission barrier, i.e. virus secretion from salivary glands.

### Expected impact of low transmission rates on capulavirus ecology

The low transmission rate of ALCV by *A. craccivora* in comparison with that of other geminiviruses is consistent with the hypothesis that the delayed discovery of capulaviruses may be ascribed to their low dissemination to and within cultivated plants. Unlike nanoviruses and luteoviruses which evolved efficient transmission with aphids, it seems that geminiviruses did not. Thus, it is thought that capulaviruses adapted to aphid transmission only in host environments where high transmission efficiency is not critical for survival. Alfalfa meets this criteria. Indeed, it is a hardy perennial plant that exhibit a high tolerance to non-biotic stresses like drought and extreme temperatures. This tolerance is expected to have at least two beneficial effects on ALCV fitness. Firstly the window of opportunity to be carried to a new plant is extremely wide and may extend over several years. Secondly, as alfalfa is hardier than other plants, it is relatively more conducive to insect feeding under biotic stresses and therefore may attract aphid vectors. The hardy perennial feature also applies to *E. caput-medusae*, the host of EcmLV, and probably also to *P. lanceolata* the host of PlLV. The french bean Severe leaf curl virus would seem to be an exception as it is the only capulavirus isolated from an annual host. However, field observations in India indicate that its incidence in French bean is generally below 2% (Akram, personal communication). Finally, according to preliminary results, ALCV did not affect the fitness of its aphid which is another factor that may have contributed to the adaptation of geminiviruses to aphid transmission.

## Conclusion

The outcome of this study, together with previous results (9), show that post-gut barriers were associated not only to the non-transmission of ALCV by a non-vector population of *A craccivora* but also to the low transmission of the vector population tested here. While such barriers found in other virus/insect combinations exhibited transmission failures (see above, TYLCCNV and PLRV), we hypothesize that the relatively low efficiency of capulavirus transmission might be constitutive of their aphid transmission adaptation. To validate this hypothesis, transmission parameters should be analyzed with other capulavirus/aphid combinations and the low transmission efficiency of capulaviruses observed in laboratory and field conditions should be further investigated.

The outcome of this study, together with previous results (9), show that post-gut barriers were associated not only to the non-transmission of ALCV by a non-vector population of *A. craccivora* but also to the low transmission of the vector population tested here. While such barriers were found with non compatible virus/insect combinations (see above, TYLCCNV and PLRV), we hypothesize that they are constitutive of the adaptation of geminivirus transmission to aphids and in turn may explain their low transmission efficiency. To validate this hypothesis, transmission parameters should be analyzed with other capulavirus/aphid combinations and the low transmission efficiency of capulaviruses observed in laboratory and field conditions should be further investigated.

## Authors and contributors

Faustine Ryckebusch (FR), Nicolas Sauvion (NS) and Michel Peterschmitt (MP) conceived and designed the experiments. FR, NS, Martine Granier and, MP performed the experiments. FR, NS, and MP analyzed the data. FR, NS, and MP wrote the paper

## Acknowledgments

We are grateful to Marie-Stéphanie Vernerey and Elodie Piroles for their assistance in the development of FISH and the use of the confocal microscope. We thank Michel Yvon, Sophie Le Blaye, Jean-Luc Macia, Sylvaine Boissinot, Véronique Brault and Romain Ferdinand for their technical assistance, Myriam Siegwaert (Inra-Avignon) for providing part of EPG material, Guillaume Sauvion for aphid drawing, and Bruno Serrate (Inra-CBGP) for the aphid picture.

## Supplementary Information

**S1 Table.**

Description of the eight transmission tests designed to estimate the transmission rate of ALCV by *A. craccivora* as a function of the number of individuals per test plants (experiment 4). The results are summarized in Fig. 4b. AAP, acquisition access period; IAP, inoculation access period; Aphid batch size, number of aphid individuals per test plant. Theoretical individual transmission rate (pi) was deduced from formula, *TR* = 1-(1-pi)^n^, knowing *TR* and n.

