## Supplemental Table 1 for "Alfalfa leaf curl virus is efficiently acquired by its aphid vector *Aphis craccivora* but inefficiently transmitted"

### Supplementary Information

**S1 Table.**

Description of the eight transmission tests designed to estimate the transmission rate of ALCV by *A. craccivora* as a function of the number of individuals per test plants (experiment 4). The results are summarized in Fig. 4b. AAP, acquisition access period; IAP, inoculation access period; Aphid batch size, number of aphid individuals per test plant. Theoretical individual transmission rate (pi) was deduced from formula , $TR = 1\text{-}(1\text{-pi})^{n}$ , knowing *TR* and n.

| | | | | **Aphid batch size (n)** | | | | | | |
|---|---|---|---|---|---|---|---|---|---|---|
| | Experiment | AAP | IAP | 1 | 5 | 10 | 20 | 30 | 40 | 100 |
| **Number of infected plants / Number of tested plants** | Test 1 | 3 | 5 | 4/43 | | | | | | |
| | Test 2 | 2 | 5 | 2/30 | 6/20 | | | | | |
| | Test 3 | 3 | 5 | 1/10 | 3/10 | 5/10 | 3/7 | | | |
| | Test 4 | 3 | 5 | 1/10 | 1/10 | 4/10 | 5/10 | 4/7 | | |
| | Test 5 | 3 | 4 | | | 4/11 | | | | 6/6 |
| | Test 6 | 4 | 5 | 1/35 | | | | | | |
| | Test 7 | 3 | 5 | 137 | | | | | | |
| | Test 8 | 3 | 2 | | 3/10 | | | | 8/10 | |
| | Total | 2-4 | 2-5 | 10/165 | 13/50 | 13/31 | 8/17 | 4/7 | 8/10 | 6/6 |

| | | | | | | | | | | |
|---|---|---|---|---|---|---|---|---|---|---|
| **% infected plants** | Test 1 | 3 | 5 | 9.30 | | | | | | |
| | Test 2 | 2 | 5 | 6.66 | 30.00 | | | | | |
| | Test 3 | 3 | 5 | 10.00 | 30.00 | 50.00 | 42.86 | | | |
| | Test 4 | 3 | 5 | 10.00 | 10.00 | 40.00 | 50.00 | 57.14 | | |
| | Test 5 | 3 | 4 | | | 36.36 | | | | 100.00 |
| | Test 6 | 4 | 5 | 2.86 | | | | | | |
| | Test 7 | 3 | 5 | 2.70 | | | | | | |
| | Test 8 | 3 | 2 | | 30.00 | | | | 80.00 | |
| | **N** | | | **6** | **4** | **3** | **2** | **1** | **1** | **1** |
| ***Transmission rate (TR)*** | **Min.** | | | **2.70** | **10.00** | **36.36** | **42.86** | | | |
| | **Max.** | | | **10.00** | **30.00** | **50.00** | **50.00** | | | |
| | **Mean** | **2-4** | **2-5** | **6.06** | **26.00** | **41.93** | **47.06** | **57.14** | **80.00** | **100.00** |
| | **SE** | | | **1.40** | **5.00** | **4.08** | **3.57** | | | |
| **Individual transmission rate (pi)** | | | | **6.1%** | **5.8%** | **5.3%** | **3.1%** | **2.8%** | **3.9%** | **-** |
